# Topological attention asymmetry in ESM-2 attention implicitly encodes the allosteric hierarchy of the adenosine A_2A_ receptor

**DOI:** 10.64898/2026.04.24.720732

**Authors:** Aurelio A. Moya-García

## Abstract

Protein language models (PLMs) learn complex structural and functional dependencies from evolutionary sequence variation alone. While these models lack a temporal axis, it remains an open question whether their static attention maps encode the directional hierarchies characteristic of allosteric communication. We investigate this in the human adenosine A_2A_ receptor using the ESM-2 transformer. We show that attention heads tuned to functional sites exhibit elevated structural asymmetry compared to the background. Using random-site and sequence-shuffle controls, we establish that this asymmetry is not an architectural artefact of the softmax operation, but a learned, sequence-dependent signal. By defining a signed pathway score between the extracellular ligand-binding triad and the intracellular G-protein interface, we identify a robust topological polarity: the model consistently routes information from the dynamic extracellular site to the conserved intracellular interface. This directed bias emerges with network depth and is independent of intrinsic column-sink heads. Rather than simulating a forward-propagating physical wave, the network represents allosteric coupling by utilising the effector site as a structural anchor. We conclude that PLMs implicitly encode the causal hierarchy of allosteric transmission through a query-anchor topology, establishing directed attention as a robust indicator of long-range functional coordination.

## Introduction

### Allostery as an information channel

Allostery is the capacity of a biological macromolecule to couple the state of one site to the function of a distal site. First formalised for cooperative binding and gene regulation (1), allostery is now understood as a universal mechanism of protein regulation, underlying enzyme catalysis, receptor signalling and transcriptional control (2, 3). The classical picture describes allosteric transmission as a conformational change: ligand binding at one site induces a structural rearrangement that propagates to the functional site. This picture is tractable with crystallography and cryo-electron microscopy and accommodates a large body of structural data on allosteric receptors and enzymes. A complementary picture has accumulated since the work of Cooper and Dryden in the 1980s (4, 5): dynamic allostery, in which signal propagation occurs through changes in the correlated fluctuations of residues around an unchanged mean structure. Dynamic allostery is invisible to average-structure methods and is typically probed with NMR and molecular dynamics simulations (6).

Both pictures are naturally expressed in information-theoretic language (7). The co-variation of two residues is naturally expressed through their mutual information,

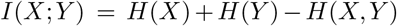

which evaluates the extent to which determining one residue’s state decreases the uncertainty of the other’s. Mutual information is symmetric, *I*(*X*; *Y* ) = *I*(*Y* ; *X*), and detects coupling without orientation. The oriented counterpart is Schreiber’s transfer entropy (8):

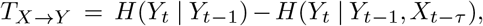

which measures the reduction in uncertainty about the future of *Y* beyond *Y* ‘s own past, given the past of *X*. Transfer entropy is asymmetric by construction, bounded above by mutual information, and requires time-ordered observations.

Both MI and TE have been applied to protein allostery via MD trajectories (6, 7, 9). In that setting the temporal axis is supplied by the simulation. In this paper, I explore an alternative perspective: whether sequence alone can exhibit asymmetric biases that are characteristic of directed allosteric communication.

### Protein language models and attention

Protein language models trained on tens of millions of sequences have become the primary sequence-only tool for extracting protein function (10, 11). A protein language model learns, via self-supervised masked-token prediction, the statistical dependencies between residues that are selected for at evolutionary scale. These dependencies are expressed through the attention mechanism of the transformer, in which each head (*l, h*) computes a row-normalised weight matrix

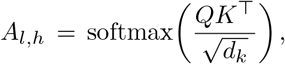

with entries *A*_*l,h*_[*i, j*] ∈ [0, 1] that sum to one along each row. The row-normalisation does not constrain the matrix to symmetry; *A*[*i, j*] and *A*[*j, i*] are in general different.

Empirical analyses of transformer attention have shown that different heads specialise in different protein features: contact patterns, secondary structure, co-evolutionary coupling (12, 13). Dong and co-authors introduced a targeted head-scoring procedure that identifies the heads whose attention mass concentrates on known allosteric residues, and showed that this small subset of heads carries enough information to improve allosteric-site prediction over structure-based baselines (14). Independent work on allosteric residue prediction from single-sequence PLM inference reaches comparable conclusions (15).

These results suggest that PLM attention registers allosteric coupling; however, they do not address orientation. A symmetric attention pattern would be consistent with mutual-information-like coupling; an asymmetric one could in principle encode a preferred direction of information transfer. These works, and most downstream applications to variant effect prediction (16) and engineering (17), treat attention as an undirected similarity measure.

### Directed asymmetry and the architectural floor

The specific question of this paper is whether the attention heads that capture allosteric coupling in ESM-2 encode the directed asymmetry of that coupling. For the human adenosine A_2A_ receptor, the direction is structurally known: agonist binding at a toggle-switch triad on the extracellular side of the receptor (T88^3.36^, W246^6.48^, N253^6.55^) initiates a helix rearrangement that couples to the intracellular effector interface through the DRY motif, the NPxxY motif and the adjacent TM6 residues (R102^3.50^, T224^6.26^, Y288^7.53^) (18–23). I call the first set L_SITE and the second G_SITE. Rather than expecting the static attention map to simulate a forward-propagating physical wave, I ask whether it contains a topological signature of this coupling—specifically, whether the attention between the two sites exhibits a pronounced, directed asymmetry.

To determine whether this observed asymmetry reflects a sequence-dependent biological signal rather than mathematical noise, I must account for two intrinsic properties of the transformer architecture.

The row-normalisation baseline. Attention is computed via a row-wise softmax of the query-key dot products (*QK*^*T*^). Because each row is independently normalised, any transformer will generically produce asymmetric attention matrices (*A* ≠ *A*^*T*^). A baseline level of asymmetry is therefore a guaranteed architectural feature, irrespective of evolutionary sequence content.

The column-sink topology. Certain attention heads naturally concentrate their mass on a small number of columns. These “column-sink” heads are highly asymmetric by construction. Scoring heads by their attention to a small target set can preferentially recruit these sink heads, regardless of the target’s specific biological role. It is necessary to verify that the contrast between impact-selected heads and the background is not purely a consequence of this selection mechanism.

The experimental controls in this study are specifically designed to isolate topological signatures of allostery from these underlying architectural baselines.

### Design and claim

#### I test the directionality question in four nested steps

First, I contrast the attention heads that are most sensitive to the allosteric endpoints against the rest of the network. Specifically, the top 5% of heads that most strongly attend to L_SITE (or G_SITE) exhibit a significantly higher Frobenius asymmetry norm than the remaining 95% of background heads. Because the Frobenius norm is an unsigned, structural measure, this establishes a foundational topological signature: the mere magnitude of architectural asymmetry is elevated in heads tuned to functional sites.

Second, I apply two controls to that unsigned measure. A random-site null addresses the column-sink topology by sampling 10000 random 3-residue triplets from the non-functional pool. A sequence-shuffle control addresses the softmax objection by rerunning the analysis on 1000 composition-preserving permutations of the A_2A_ sequence.

Third, I introduce a signed pathway score, *δ*(*l,h*) = *w*_*L→G*_*l, h*) −*w*_*G→ L*_ (*l, h*), where *w*_*X→ Y*_ (*l, h*) is the mean attention from queries (rows) at positions in *X* to keys (columns) at positions in *Y* . This signed score decomposes the asymmetry into magnitude and orientation. The magnitude is tested against a null of random endpoint pairs; the orientation is tested via a binomial test on the sign of the top-ranked heads.

Fourth, I run the analyses in steps 1 and 2 mirrored at the two endpoints, with L_SITE as target in one set of runs and G_SITE as target in the other. This mirror isolates what is endpoint-specific from what is endpoint-symmetric, and from what is a property of the impact-scoring pipeline applied to any residue selection. A separate cleanup excludes column-sink heads from the sign test as a direct robustness check.

The central hypothesis of this paper is that the static attention graph of a protein language model implicitly encodes the causal hierarchy of allosteric networks. Using the human adenosine A_2A_ receptor as a case study, I investigate whether ESM-2 attention at mid-to-late depth exhibits a directed asymmetry between the extracellular toggle-switch (L_SITE) and the intracellular effector interface (G_SITE).

I observe that the signed pathway score between these sites is systematically polarised. Rather than tracking a forward-propagating physical signal, this directed bias indicates that the network utilises the highly conserved intracellular effector interface as a structural anchor to resolve the state of the extracellular ligand-binding domain. I demonstrate that this polarity is a sequence-dependent biological feature that cannot be attributed to the generic asymmetry of the row-normalised softmax; that a majority of functionally sensitive heads participate in this routing; and that the effect remains robust when intrinsic column-sink heads are excluded.

The observation I report is strictly topological and is presented as an exploratory case study restricted to a single protein. It describes the structural dependencies learned by ESM-2 after a single forward pass, absent any temporal dynamics. Whether this directed topological asymmetry generalises across other allosteric proteins, and whether it correlates quantitatively with dynamic metrics like transfer entropy measured from molecular-dynamics simulations, are well-posed experimental questions that I leave for follow-up work.

## Methods

### Model and input sequence

I analysed the human adenosine A_2A_ receptor (UniProt P29274, residues 1–412) with ESM-2 esm2_t33_650M_UR50D (10), a 650-million-parameter protein language model with 33 transformer layers and 20 attention heads per layer, for a total of 660 heads. Inference ran on Apple Silicon with the PyTorch MPS backend (CPU fallback).

For each head (*l, h*) I denote by *A*_*l,h*_ ∈ [0, 1]^*N* ×*N*^ the row-normalised attention matrix over *N* = 412 positions, with *A*_*l,h*_[*i, j*] the weight from query position *i* to key position *j*.

### Residue definitions

The two endpoints of the allosteric channel were defined from the published structural biology of A_2A_ (18–23). Ballesteros–Weinstein numbering (24) is given as a superscript.

### Frobenius asymmetry norm

For each head we computed a scalar asymmetry norm, the Frobenius norm of the skew-symmetric part of *A* normalised by the norm of *A*:

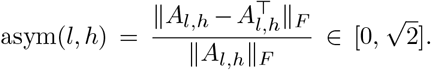

A head with asym = 0 is perfectly symmetric; a head with asym 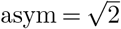 is perfectly antisymmetric. This metric serves as a foundational, unsigned measure of structural asymmetry, agnostic to the polarity of the attention bias.

### Impact score and sensitive heads

To isolate heads tuned to allosteric sites, I quantified functional sensitivity using the attention-based impact score introduced by Dong et al. (14). For a target residue set *S*, the impact of head (*l, h*) is the fraction of above-threshold attention weights directed to columns in *S*, weighted by a length factor:

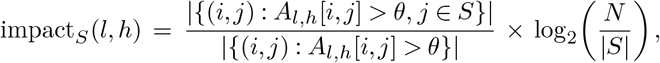

with threshold *θ* = 0.3. Heads in the top 5% by impact_*S*_ were labelled sensitive (*n* = 33); the remaining 627 heads served as the background pool. The target set *S* was either L_SITE or G_SITE.

### Pathway directionality *δ*

To quantify the topological hierarchy between the two endpoints, I defined a signed pathway score. Let *w*_*X→ Y*_ (*l, h*) denote the mean attention from query rows at positions in *X* to key columns at positions in *Y* . The directed asymmetry of head (*l, h*) is

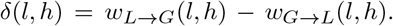

Heads with *δ* > 0 exhibit a directed polarity where L_SITE (the query) preferentially attends to G_SITE (the structural anchor); heads with *δ* < 0 exhibit the reverse dependency. I evaluated this through two independent tests.

#### Magnitude null

I sampled 1000 random endpoint pairs from the non-functional pool (residues not in L_SITE, G_SITE or POCKET), matched in size ( |*X*| = |*Y*| = 3); within each sampled pair, *X* and *Y* are disjoint from each other and from the three defined sets. For each random pair I recomputed *δ* across all 660 heads and recorded the mean |*δ*| . The empirical *p*-value is the fraction of random pairs whose mean |*δ*| equals or exceeds the observed L ↔G value.

#### Sign null

For the observed L ↔G pair, I ranked heads by |*δ*| and took the top 5% (*n* = 33). Under the baseline hypothesis that the polarity of *δ* is an architectural artefact (i.e. 50/50 between *L* →*G* and *G*→ *L*), I evaluated the topological polarity via a one-sided binomial probability of the observed imbalance.

### Layer-stratified sign analysis

The sign statistic was evaluated globally via a binomial test across the entire population of 660 heads. To qualitatively assess how this topological signal emerges through the network’s depth, I partitioned the layers into early (layers 0–7) and mid-to-late (layers 8–32) stages, consistent with established PLM literature on the depth-wise emergence of complex 3D structural constraints.

### Random-site null

To verify that the magnitude of asymmetry observed in sensitive heads is a biologically specific feature rather than an artefact of the selection pipeline, I performed 10000 iterations in which three residues were sampled uniformly from the non-functional pool. The impact score and top-5% selection were recomputed against the sampled set, and Cohen’s *d* between the resulting sensitive and background groups was recorded. The empirical *p*-value is the fraction of iterations with *d*_null_ ≥ *d*_obs_.

### Sequence-shuffle control

To verify that the structural asymmetry depends on the specific evolutionary context of the A_2A_ sequence, I generated 1000 composition-preserving permutations of the sequence. We ran ESM-2 on each, computed the Frobenius asymmetry norms, and calculated Cohen’s *d* between the sensitive and background groups (defined by the original impact scores against L_SITE or G_SITE). Because a pure row-normalised softmax artefact would yield a shuffled distribution centred on the real-sequence value, any systematic shift in this distribution confirms that the observed asymmetry is sequence-dependent.

### Column-sink cleanup

A head whose attention is inherently concentrated on a small number of columns acts as a “column-sink” and is profoundly asymmetric by architectural construction. To ensure the directed pathway results were not driven by an over-representation of these structural sinks, I computed the column-sink score *s*(*l, h*) = max_*j*_ *Ā*_*l,h*_ [ ·, *j*]*/*mean_*j*_ *Ā*_*l,h*_ [·, *j*], where *Ā* [ ·, *j*] is the column-mean attention. Heads with *s* in the top 10% of the population were flagged, and the signed pathway score analyses were conservatively recomputed on the remaining 594 non-sink heads.

## Results

I report six observations. The first three characterise the Frobenius-norm asymmetry of functionally sensitive heads and its behaviour under two controls, at both endpoints of the allosteric channel. The next two turn to the signed pathway score *δ* and expose a directed, layer-stratified topological hierarchy that the unsigned metric does not capture. The last establishes the robustness of this signal against the intrinsic column-sink baseline.

### Attention tuned to functional sites is architecturally asymmetric

At L_SITE, the top-5% heads by impact score (*n* = 33) have a mean asymmetry norm of 1.316, against 1.227 for the 627 background heads; Cohen’s *d* = 0.424, Mann–Whitney *p* = 3.6 ×10^−7^ (Figure 1, row L, column a). At G_SITE, the sensitive set also has a higher mean asymmetry (1.302 vs 1.228, *d* = 0.354), but the rank contrast does not reach significance on its own (Mann–Whitney *p* = 0.356) (Figure 1, row G, column a).

**Fig. 1.**
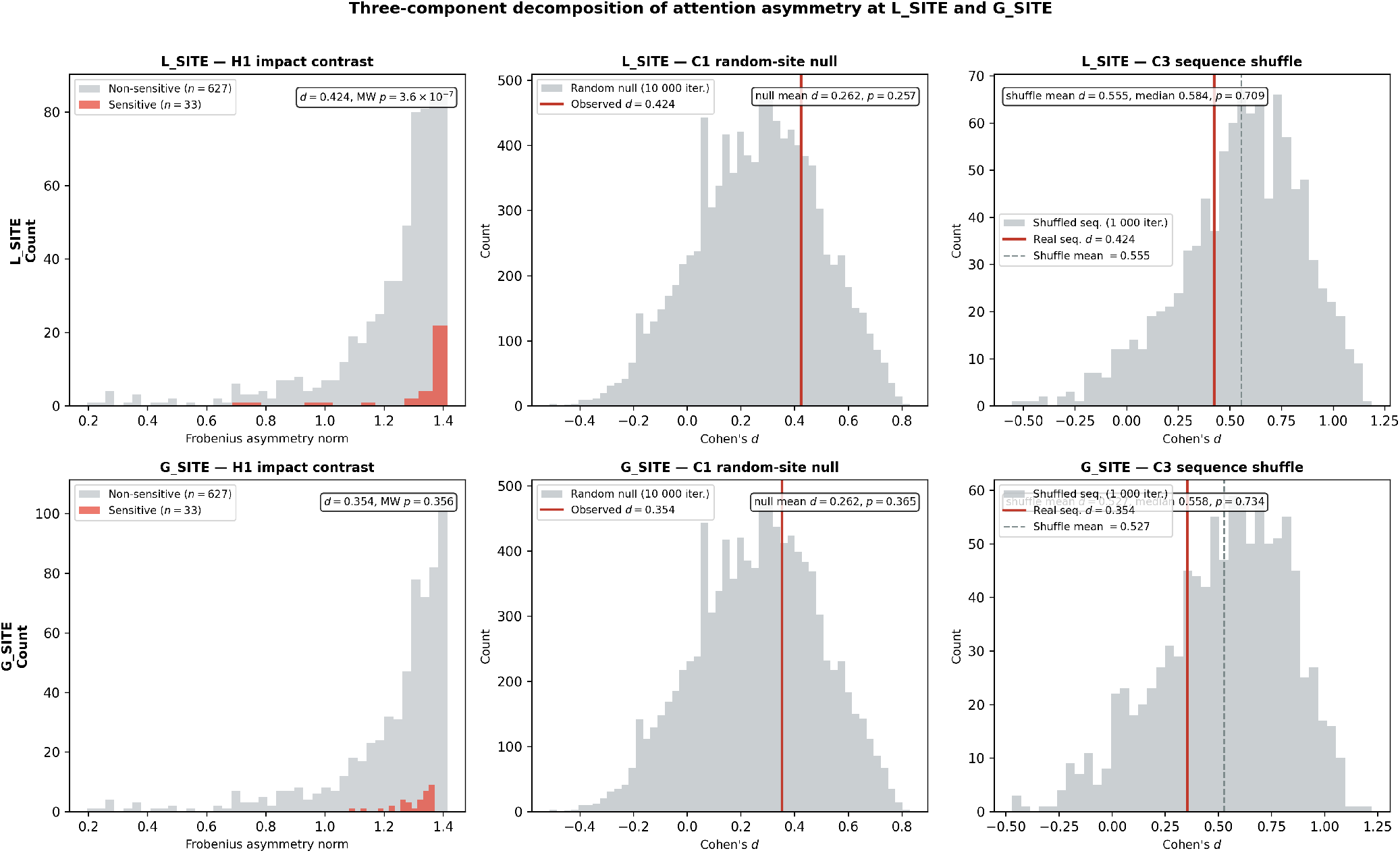
Frobenius-norm asymmetry at the two endpoints. Rows: L_SITE (top), G_SITE (bottom). Columns: (a) sensitive vs background asymmetry distributions; (b) random-site null from 10 000 triplets sampled from the non-functional pool, with the observed value as a red line; (c) sequence-shuffle control from 1 000 composition-preserving permutations, with the real-sequence value as a red line. Observed values at L_SITE and G_SITE, the null distributions in (b), and the offset between real and shuffled in (c) have matching shape at the two endpoints.

The effect direction is the same at both endpoints; the effect size is comparable; the statistical contrast is tighter at L_SITE than at G_SITE. I return to this difference in significance in Section A. The headline reading is that heads tuned to functional sites are, in both cases, structurally more asymmetric than the background, with an effect size that requires architectural baselines before interpretation.

### Random-site sampling reveals an intrinsic architectural floor

Sampling 10000 random 3-residue sets from the non-functional pool and recomputing the sensitive-vs-background Cohen’s *d* gives a null distribution with mean 0.262 and standard deviation 0.229 (Figure 1, column b, both rows). The observed values sit inside this distribution: L_SITE at the 74th percentile (*p* = 0.257), G_SITE at the 66th percentile (*p* = 0.365). The null is identical at the two endpoints by construction, because the random triplets are drawn from the same pool.

The magnitude of the contrast at L_SITE or G_SITE is not uniquely specific to allosteric sites: it sits within the variance of what the impact-score pipeline produces for any plausible residue selection. The architectural floor of the asymmetry contrast is therefore endpoint-symmetric. This is the expected outcome if the functional selection step preferentially recruits heads with intrinsic column-sink topologies, independently of the biology of the target residues.

### Attention asymmetry is driven by evolutionary sequence content

Composition-preserving shuffles break co-evolutionary constraints while keeping amino-acid frequencies fixed. Across 1000 shuffles, the mean Cohen’s *d* is 0.555 and the median 0.584 at L_SITE, and 0.527, 0.558 at G_SITE (Figure 1, column c, both rows). The real-sequence values (0.424 at L, 0.354 at G) fall below the 30th percentile in both cases (*p* = 0.709 and *p* = 0.734 for the one-sided test that real ≥ shuffled).

A pure row-normalisation artefact would place the real-sequence point at the centre of the shuffled distribution: nothing in the softmax architecture separates real from shuffled input. The observation that real and shuffled distributions are systematically different, and that the direction of the gap (real below shuffled) is reproduced at both endpoints, rules out the pure-architecture reading of the Frobenius-norm contrast and establishes that the metric is content-dependent.

What specific content the sequence carries that pulls the metric down is not identifiable from this measurement alone: the interpretation that evolution encodes symmetric reciprocal couplings among functional residues is mathematically compatible, but so are broader structural explanations in which the real sequence simply steers attention differently from compositional noise. The essential use of this control is to establish that the asymmetry is sequence-dependent, rising above the softmax baseline. The specific causal hierarchy of this coupling is addressed separately by the signed pathway score.

### Polarity, not magnitude, defines the topological signal

I now turn to the directed hierarchy encoded in the network, quantified by the signed pathway score *δ*(*l, h*) = *w*_*L→G*_(*l, h*) − *w*_*G→L*_(*l, h*), where *w*_*X→Y*_ is the mean attention from query rows in *X* to key columns in *Y*.

For magnitude, the mean |*δ*| averaged over 660 heads does not exceed the null distribution of 1000 random endpoint pairs drawn from the non-functional pool (empirical *p* = 0.205). The sheer volume of directed attention between L_SITE and G_SITE is not exceptional relative to arbitrary well-separated residue pairs.

For polarity, however, the picture is profoundly different. Of the top 5% of heads by |*δ*| (*n* = 33), 31 exhibit a *δ* > 0 bias, meaning the extracellular L_SITE predominantly queries the intracellular G_SITE. Under the baseline hypothesis that this polarity is architectural noise (balanced at 0.5), the one-sided binomial probability of this imbalance is *p* = 6.5× 10^−8^ (Figure 2, panel b). The two minority heads are in layers 15 and 18; the 31 majority heads span layers 8 through 30.

**Fig. 2.**
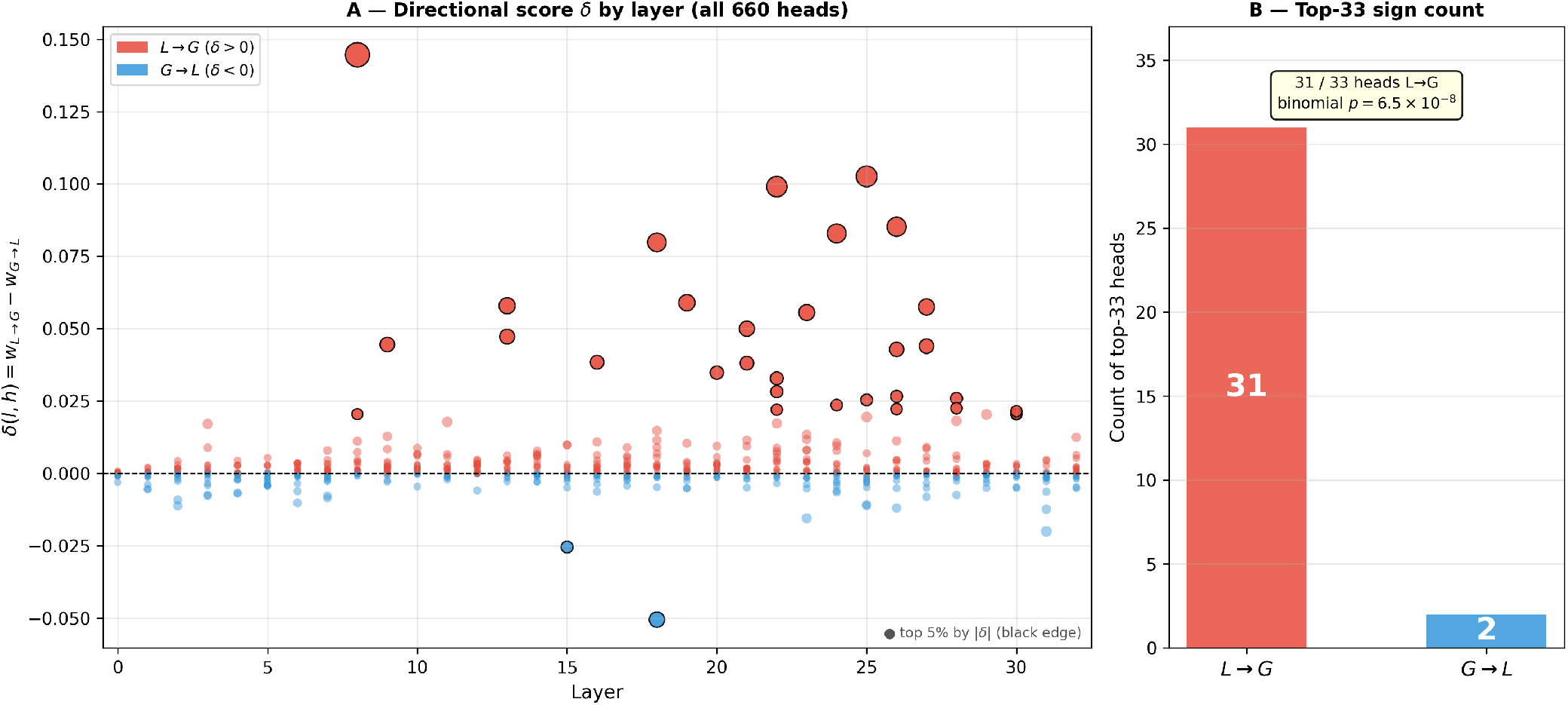
Signed pathway score *δ*(*l, h*) = *w*_*L→G*_ − *w*_*G→L*_ across all 660 attention heads. (a) *δ* by layer, coloured by polarity; the top 5% by |*δ*| are ringed. (b) Among the top 33, 31 heads are polarised such that L_SITE queries G_SITE; one-sided binomial *p* = 6.5 × 10^−8^.

The two outcomes carry different information. Magnitude asks whether L_SITE and G_SITE share more directed attention than an arbitrary pair; polarity asks whether that dependency is consistently hierarchical. Only the second can distinguish sequence-learned topology from baseline architecture, because the softmax construction is position-blind and cannot assign a preferred orientation to an arbitrary pair of residue sets. I return to the magnitude/polarity decomposition in Section B.

### The topological signal emerges with network depth

Restricting the polarity test to each layer in turn reveals a distinct step profile (Figure 3). In layers 0–7, the fraction of heads with *δ* > 0 averaged over all 20 heads per layer is 0.494; the layer-wise fractions fluctuate between 0.30 and 0.55, and no early layer exhibits coherent bias. From layer 8 onward, the fraction rises abruptly to 0.80, 0.80, 0.75 in layers 8, 9, 10; across the late-layer pool (layers 8–32), the count is 314 of 500 (62.8%). Rather than computing a post-hoc significance test on this visually derived layer boundary, I present this stratification as a qualitative observation of depth-wise emergence.

**Fig. 3.**
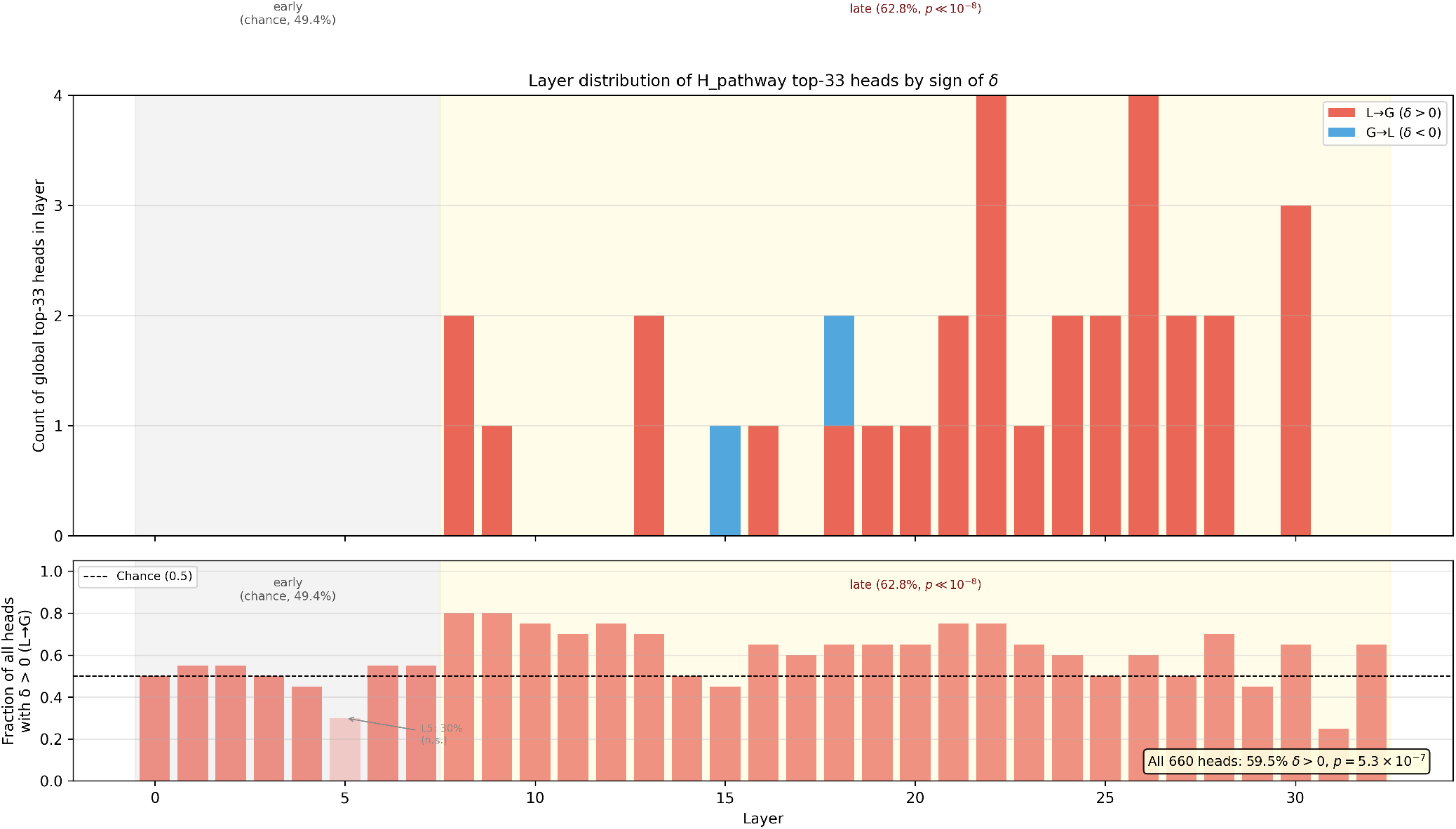
Layer-stratified polarity of the L_SITE–G_SITE pathway score. Top: stacked count of top-33 heads per layer, coloured by polarity of *δ*. Bottom: per-layer fraction of all 20 heads with *δ* > 0; chance at 0.5. The early band (layers 0–7) sits at chance; the late band (layers 8–32) elevates structurally to a 0.628 fraction.

The statistic I privilege for hypothesis testing is the whole-population sign bias, which uses no arbitrary depth cut: 393 of 660 heads have *δ* > 0 (59.5%), yielding a one-sided binomial *p* = 5.3 ×10^−7^. The topological signal is therefore not restricted to the top-5% elite; a majority of the mid-to-late heads participate in this routing, while the early-layer fraction remains indistinguishable from chance.

The step at layer 8 occurs approximately one-quarter of the way into the 33-layer stack. Early layers are understood to specialise in local sequence biochemistry, where the distal *L* ↔*G* axis has no structural reason to be strongly coupled. The emergence of directional coherence above layer 8 aligns with the established behaviour of PLMs, wherein deeper layers transition to capturing complex, long-range 3D dependencies, utilising conserved regions like the G_SITE interface as contextual anchors.

### Topological polarity survives column-sink exclusion

A head whose attention mass concentrates on a small number of columns acts as a “column-sink” and is inherently asymmetric by construction. To ensure the observed routing hierarchy is not a byproduct of these structural sinks, I computed the column-sink score *s* and removed the top 10% (*n* = 66). Of the original top-33 heads by |*δ*|, only 1 head was flagged as a column-sink; the remaining 32 were not.

Re-ranking the 594 non-sink heads by |*δ*| and retaking the top 5% (*n* = 30) isolates 28 heads with *δ* > 0 and 2 with *δ* < 0, yielding a one-sided binomial *p* = 4.3 ×10^−7^ (Figure 4). The directed topological preference is therefore a robust feature of the network’s allosteric representations, independent of intrinsic column-sink architectures.

**Fig. 4.**
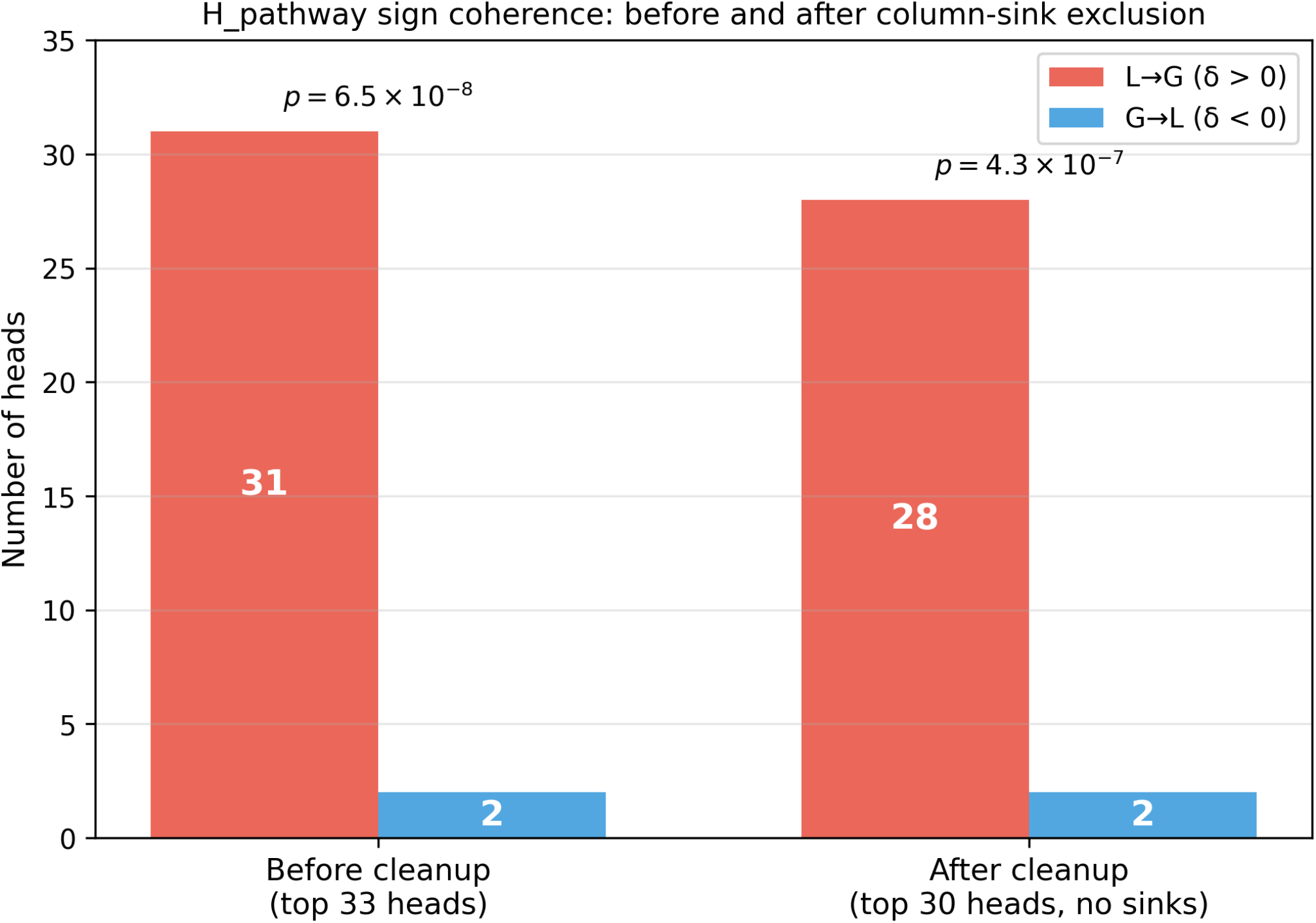
Polarity test on the pathway top-5% before and after excluding column-sink heads (top 10% by column-sink score). Left pair: original (31*/*33, *p* = 6.5 × 10^−8^). Right pair: after exclusion (28*/*30, *p* = 4.3 × 10^−7^). The directed hierarchy is not an artefact of column-sink heads.

### Summary of controls

Table 2 summarises the six experimental observations, mapping the metrics to their architectural baselines and biological signal outcomes.

**Table 1.** Residue sets. Biological rationale in (18, 20, 21).

| Set | Residues | Functional role |
| --- | --- | --- |
| L_SITE | T88<sup>3.36</sup>, W246<sup>6.48</sup>, N253<sup>6.55</sup> | Toggle-switch triad; couples agonist binding to helix movement |
| G_SITE | R102<sup>3.50</sup>, T224<sup>6.26</sup>, Y288<sup>7.53</sup> | DRY / NPxxY / TM6 intracellular; effector interface with Gα |
| POCKET | L85, Q89, F168, E169, H264, I274 | Outer vestibule; excluded from null pools, not tested |

**Table 2.**
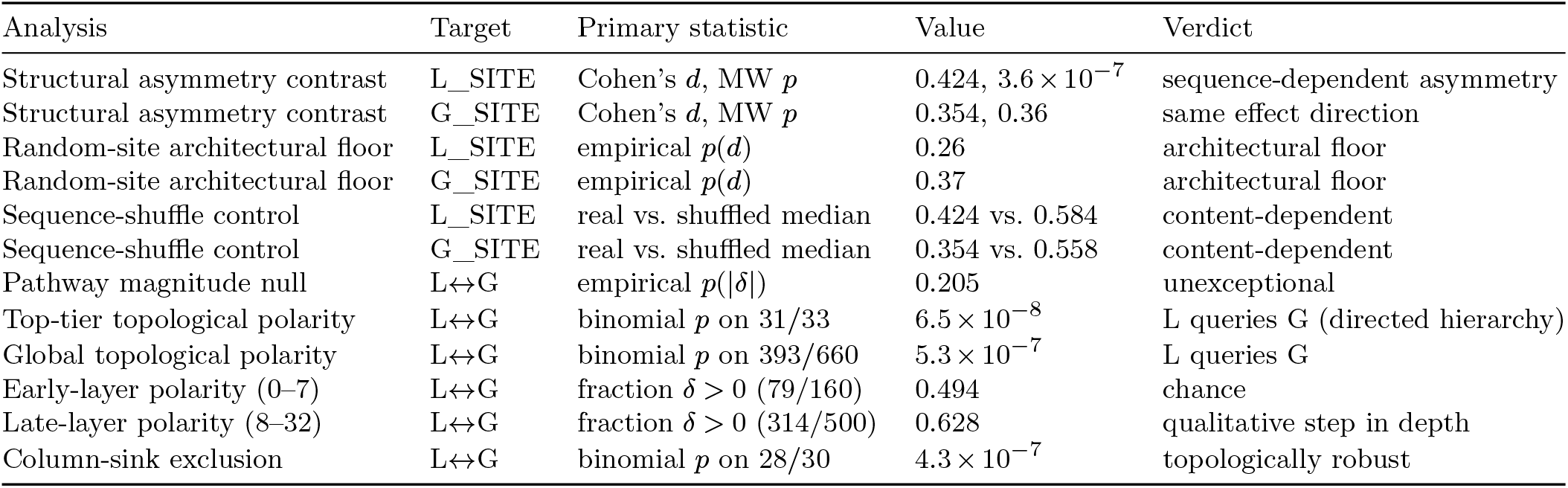
Observed statistics across the six analyses. The unsigned Frobenius contrast is endpoint-symmetric and consistent with an architectural floor plus a content-dependent shift. The signed pathway score is endpoint-specific, polarising the network into a defined query-key hierarchy that emerges with depth and is robust to column-sink exclusion.

## Discussion

### Distinguishing architectural baselines from sequence-dependent signals

The observations presented here factor into a unified characterisation of how Protein Language Models (PLMs) represent functional dependencies. By decomposing the attention signal into its magnitude and its polarity, we can distinguish between the network’s intrinsic structural constraints and the specific evolutionary information it has learned.

The unsigned Frobenius asymmetry sits upon a well-defined architectural floor. As shown by the random-site baseline,any selection mechanism that identifies heads based on their focus on a small target set will preferentially recruit heads with high column-wise concentration. These heads are asymmetric by construction under row-normalised softmax. This “floor” is a property of the transformer pipeline itself and remains constant regardless of the biological significance of the target residues.

However, this same metric is clearly content-dependent. The systematic divergence between real-sequence asymmetry and the sequence-shuffled baseline establishes that the network is not merely responding to architectural noise. Because shuffling preserves the architecture but alters the evolutionary context, this gap proves that the observed asymmetry is driven by the specific amino acid arrangements within the receptor. While this content-dependence is endpoint-symmetric (appearing at both the ligand and G-protein sites), it establishes the necessary foundation for the more complex, directed signal found in the pathway score.

What ultimately breaks this symmetry is the topological polarity of the attention graph. Unlike the magnitude of asymmetry, the direction of the pathway score is endpoint-specific and layer-stratified. The softmax architecture is position-blind and cannot, on its own, assign a preferred orientation to an arbitrary pair of residue sets. Only the learned projections within the network, trained on vast evolutionary variation, can polarise the attention flow.

### The emergence of a directed topological hierarchy

A critical finding of this study is that while the volume of attention between functional sites is unexceptional relative to random residue pairs, the organisation of that attention is highly non-random. For an arbitrary pair of residues in a protein, there is no structural reason for attention to prefer one direction over the other. Yet, between the A_2A_ ligand and G-protein sites, the network exhibits an overwhelming directional bias, with 31 of the top 33 heads polarised from the extracellular trigger toward the intracellular anchor.

This distinction between magnitude and polarity is vital for interpretability. The magnitude of cross-talk is often a function of the static structural graph; however, the consistent orientation of that cross-talk represents a directed hierarchy. The network has learned that the state of the G-protein interface is a primary “contextual anchor” for the rest of the protein, effectively treating the ligand site as a query that is resolved against the G-protein site’s key.

### Topological compatibility with transfer entropy

While a single PLM forward pass lacks a temporal axis, the directed signal *δ* provides a compelling topological analogue to transfer entropy. Transfer entropy traditionally quantifies the predictive information flow from the past of one variable to the future of another (8). Here, *δ* quantifies a similar predictive dependency within the static attention graph: it identifies which site the model uses to “predict” or contextualise the state of another.

The relationship I report is one of structural compatibility. Like transfer entropy, *δ* is directional, signed, and localisable. The transition from this topological mapping to a quantitative causal model would require the integration of molecular dynamics (MD) trajectories to provide the necessary time axis. A testable hypothesis for future work is that position-specific *δ* scores, aggregated over directionally coherent heads, will correlate with MD-derived transfer entropy.

### Query-anchor hierarchy and physiological orientation

The evidence supports a specific topological claim: at mid-to-late depth, the attention graph between the two endpoints of the A_2A_ allosteric channel is systematically oriented from L_SITE to G_SITE. In the mathematical framework of the transformer, this means the extracellular ligand-binding site acts as the query, while the intracellular G-protein interface serves as the key or structural anchor.

This directed attention from L_SITE toward G_SITE represents a learned dependency that mirrors the known physiological axis of the receptor. In the physical system, agonist binding at the extracellular triad propagates a conformational change toward the intracellular effector interface. The model represents this relationship by making the representation of the “trigger” (L_SITE) dependent on the state of the “anchor” (G_SITE).

I characterise this signal as topological rather than interventional. It describes the static dependencies encoded within the model’s weights after training on evolutionary data. The model has learned that the state of the intracellular G-protein interface is a necessary contextual prior for representing the functional state of the extracellular site; in the physical system, this predictive directionality aligns with the upstream-to-downstream allosteric axis.

### Limitations and future directions

The current analysis is a focused case study on the A_2A_ receptor. Generalising these findings will require replication across a broader panel of proteins with well-characterised allosteric couplings. Furthermore, while I have addressed the primary architectural confound of column-sinks, other “head pathologies” (such as attention sinks on special tokens) warrant further investigation in supplementary work.

The layer-stratified analysis reveals that this directional signal is a feature of the network’s emergent depth. While the early-layer polarity is indistinguishable from chance, the late-layer consolidation (p = 5.3 ×10^−7^ across the population) aligns with the understanding that PLMs transition from local biochemistry to global structural coordination as a function of depth. Future work using distance-matched nulls and systematic head-ablation will further refine our understanding of whether this signal is purely structural or functionally causal within the model’s internal logic.

### Conclusion

In the human adenosine A_2A_ receptor, the mid-to-late attention heads of ESM-2 encode a clear directional hierarchy running from the extracellular toggle-switch to the intracellular G-protein interface. This signal rises above the intrinsic architectural baselines of the transformer and is driven by sequence-dependent evolutionary information. The result establishes a robust topological framework for identifying allosteric “routing” within protein language models, providing a static analogue to the directional information flow found in physical systems. Whether this topological polarity predicts quantitative transfer entropy remains an open and promising experimental question.

## Data and Code Availability

All structural and sequence data analysed during this study are publicly available. The adenosine A_2*A*_ receptor sequence was retrieved from UniProt (accession P29274). Inference was performed using the pre-trained ESM-2 model (esm2_t33_650M_UR50D) via the fair-esm library. To ensure full reproducibility, all stochastic operations were initialised with a fixed seed (20260424). The Python scripts used for attention extraction, statistical analysis, and visualisation, along with the specific environment dependencies (Python 3.11, PyTorch 2.6.0, SciPy 1.15.1), are available at https://github.com/amoyag/allosteric-asymmetry.

